# Optimal response to quorum-sensing signals varies in different host environments with different pathogen group size

**DOI:** 10.1101/775478

**Authors:** Liqin Zhou, Leyla Slamti, Didier Lereclus, Ben Raymond

**Affiliations:** School of Biological Sciences, Royal Holloway University of London, Egham, TW20 0EX, UK; Micalis Institute, INRA, AgroParisTech, Université Paris-Saclay, 78350 Jouy-en-Josas, France; Imperial College London, Silwood Park campus, Ascot, SL5 7PY, UK; University of Exeter, Penryn campus, Penryn, TR10 9FE, UK

**Keywords:** cheating, cooperation, PlcR / PapR, polymorphism, signalling, virulence

## Abstract

The persistence of genetic variation in master regulators of gene expression, such as quorum-sensing systems, is hard to explain. Here, we investigated two alternative hypotheses for the prevalence of polymorphic quorum-sensing in Gram-positive bacteria, *i.e*. the use of different signal / receptor pairs (‘pherotypes’) to regulate the same functions. First, social interactions between pherotypes or ‘facultative cheating’ may favour rare variants that exploit the signals of others. Second, different pherotypes may increase fitness in different environments. We evaluated these hypotheses in the invertebrate pathogen *Bacillus thuringiensis*, using three pherotypes expressed in a common genetic background. Facultative cheating occurred in homogenized hosts, in contrast, rare pherotypes had reduced fitness in naturalistic infections. There was clear support for environment-dependent fitness: pherotypes varied in responsiveness to signals and in mean competitive fitness. Notably, competitive fitness varied with group size: the pherotype with highest responsiveness to signals performed best in smaller hosts where infections have a lower pathogen group size. Less responsive pherotypes performed best in larger hosts. Results using homogenized insect media fit with the expectation of facultative cheating and social evolution theory, but results from naturalist oral infections do not fit many of the predictions from this body of theory. In this system, low signal abundance appears to limit fitness in hosts while the optimal level of response to signals varies in different host environments.

**Importance:** Quorum sensing describes the ability of microbes to alter gene regulation according to their local population size. Some successful theory suggests that this is a form of cooperation: investment in shared products is only worthwhile if there are sufficient bacteria making the same product. This theory can explain the genetic diversity in these signaling systems in Gram-positive bacteria such as *Bacillus* and *Staphylococcus*. The possible advantages gained by rare genotypes (which can exploit the products of their more common neighbours) could explain why different genotypes can coexist. We show that while these social interactions can occur in simple laboratory experiments they do not occur in naturalistic infections using an invertabrate pathogen, *Bacillus thuringiensis*. Instead our results suggest that different genotypes are adapted to different-sized hosts. Overall, social models are not easily applied to this system implying that a new explanation for this form of quorum sensing is required.

## Introduction

Pathogenic bacteria often coordinate the secretion of a wide range of virulence factors via ‘quorum-sensing’ (QS), the release of -and response to-small diffusible auto-inducer molecules [1, 2]. Since the concentration of auto-inducers in the environment increases with cell density, these molecules can drive density and growth phase dependent changes in gene expression [1, 2]. The evolutionary biology of quorum sensing has been closely linked to theories of cooperation. QS signalling molecules can behave as ‘public goods’ [3, 4], products or services that benefit entire communities but which require costly contributions from individuals. In addition, QS systems may regulate the expression of virulence factors that are also public goods [3, 4]. Autoinducer molecules, especially oligopeptides, are metabolically costly [5]. Non-producers can behave as ‘cheats’: in mixed cultures they gain a growth advantage by avoiding costs of signalling and exploiting the signals produced by others [3, 4, 6–8]. Importantly, if QS regulates the production of typical public goods, theory predicts increased levels of public goods should increased group level benefits, typically total bacterial population size [4, 9–11]. However, in pathogens such as *Bacillus thuringiensis*, virulence factors do not typically behave as classical public goods and confer increased mortality or increased invasiveness as benefits [12, 13].

QS systems are diverse both within and between species. Gram negative bacteria often have systems based on acyl-homoserine lactones while Gram positive bacteria commonly have peptide based auto-inducing signals [2, 14]. Many species have multiple QS systems with different mechanisms activating different genes [15]. However, in many peptide-based systems in *Bacillus, Streptococcus* and *Staphylococcus*, there is unexplained genetic diversity in quorum-sensing signals and receptors that activate the same presumptive suite of genes [16–20]. Different QS alleles can be found in the same ecological niche and within closely related clades [21–23]. The functional significance of these polymorphic signal variants, known as pherotypes, is largely unknown.

In the *Bacillus cereus* group, PlcR is a master regulator of virulence, which controls the production of proteins involved in overcoming host barriers, host immune defence and bacterial competition [24]. The function of PlcR is activated by the binding of the signalling peptide PapR [25]. There are four distinct pherotypes in the PlcR/PapR system [20, 23]. Each pherotype (Groups I-IV) corresponds to a distinct signalling peptide: PapRI A(G/S/K)DLPFEF(Y); PapRII SDMPFEF, PapRIII N(S/Q)E(D)VPF(Y)EF(Y), PapRIV SDLPFEH, but these pherotype alleles are not restricted to particular clades or species [20]. For instance, all four pherotype alleles have been found in *B. thuringiensis* [20], a specialized invertebrate pathogen and one of the common species in the *B. cereus* group [26]. There are varying degrees of ‘cross-talk’ between pherotypes in this system, this being the activation of one receptor by the signal from another pherotype. Receptor-peptide interactions can be highly specific, in Group IV for example, while Group II and III receptors can be activated to a lesser extent by peptides from other Groups [23] (Fig 1). PlcR receptor proteins vary in the region involved in signal binding but have highly conserved helix-turn-helix domains, the region involved in DNA binding [23, 27]; different PlcR pherotypes are expected to activate the same suite of genes. Thus the persistence of diverse pherotypes remains a puzzle.

**Fig 1.**
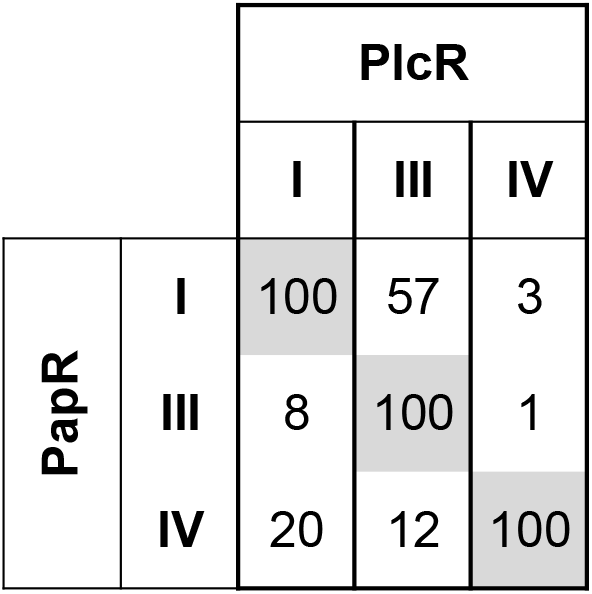
Variation in responsiveness and in cross-talk between QS pherotypes in the *B. cereus* group. Each column presents the % of transcriptional activity of P_*plcA*_, a PlcR-dependent promoter, when the indicated peptide is added to the cells relative to its activity when the cognate peptide is used for each PlcR pherotype. Each column should be read independently of the others. PlcR I, III, IV indicate the *plcR* allele harbored by an otherwise isogenic *ΔplcRpapR* strain. PapR I, III, IV indicate the pherotype of the synthetic peptide added to the cells. The figure was produced using data from Bouillaut et al. (2008).

One evolutionary phenomenon that can maintain high levels of polymorphism is negative frequency dependence, the concept that rare variants can have a fitness advantage over more common types. Social interactions (cooperation / cheating) between QS variants can provide a mechanism for negative frequency dependence: QS cheaters or non-producers commonly have higher fitness when rare in mixed cultures [3, 4, 6–8]. One form of social exploitation that could drive maintenance of polymorphism is ‘facultative cheating’ [8, 28, 29]. Facultative cheating can occur if pherotypes respond less efficiently to signals from bacteria with different QS alleles, *i.e*. if cross-talk between variants is limited. Rare pherotypes may act as transient cheats, since at low frequencies they will have low-level responses to their own QS signals, but may still exploit the secreted products of the common pherotypes. Facultative cheating can drive frequency dependent cheating behaviour and negative frequency dependent fitness [8, 28].

An alternative hypothesis for QS polymorphism is that different pherotypes have varying fitness in distinct environments [16, 30]. Importantly, for the *B. cereus* group, different pherotypes vary in the degree of responsiveness to their own peptide signal peptides: standard concentrations of signal produce different levels of transcription of a PlcR dependent reporter [23]. The structural basis of the interactions between PapR and PlcR leading to the activation of PlcR has been reported for pherotype I [27, 31], and structural modelling has provided insight into the binding of PapR II, III or IV to their corresponding PlcR variant (Bouillaut 2008). Other things being equal, more responsive pherotypes are expected to invest more in virulence factors than less responsive counterparts. One important environmental variable predicting investment in QS dependent virulence is group size: increased investment in cooperative virulence should be favoured in larger groups [9].

We tested both the facultative cheating and environment-dependent fitness hypotheses experimentally, using different pherotypes from diverse *B. thuringiensis* strains expressed in a common genetic background. These hypotheses make distinct predictions: the facultative cheating hypothesis predicts that alleles will have higher fitness when rare in mixed infections (negative frequency dependence) and assumes that all QS are phenotypically equivalent, while the environment dependent hypothesis predicts that distinct alleles produce distinct phenotypes which vary in fitness according to environmental conditions. In addition, we tested the environment dependent fitness hypothesis by characterizing the phenotypic differences conferred by different pherotype alleles. Since early results indicated that the relative fitness of competing pherotypes varied with abundance of *B. thuringiensis* in oral infections, we also tested how factors that determine pathogen group size (i.e. size of host and pathogen dose) affect competitive fitness.

While there is support for facultative cheating from simple *in vitro* systems, this mechanism requires relatively homogeneous, well-mixed conditions where co-existing genotypes can exploit each other’s secreted products [8, 28, 32]. Using the same insect host and set of mutants we tested the facultative cheating and environment dependent hypotheses in two different experimental regimes where spatial structure and opportunities for exploitation of QS vary considerably. In the first regime, which uses homogenized insects as a growth medium, PlcR and PapR non-producers can behave as cheats and the benefit provided by QS is increased final population size [12], in common with many classic bacterial public goods [3, 4, 11]. The second experimental regime uses oral infections of *B. thuringiensis*, the natural mode of infection [33]. In this regime, there is strong spatial structure, PlcR and PapR non-producers are not effective cheats, while the key benefits of QS arise from increased infectivity and improved ability to invade host tissues from the midgut [12, 25, 34].

## Results

### Differences in expression of QS-regulated virulence factors between pherotypes

In order to determine if there was a variation in PlcR activity between pherotypes, we measured the production of QS regulated lecithinase *in vitro* via hydrolysis of egg yolk phosphatidylcholine. We also conducted assays of QS-dependent transcription rate, measured by a β-galactosidase reporter. Peak (maximal) enzyme activity for lecithinase and ß-galactosidase occurred one and two hours after entry into the transition phase between exponential growth and stationary phase and was consistent across all pherotypes (data not shown). The Group IV pherotype had the highest peak lecithinase activity (Fig 2A; square-root transformed data, Likelihood ratio test = 7.04, *df* = 2, *P* = 0.0295), while the other two pherotypes were indistinguishable from each other (Likelihood ratio test = 0.61, *df* = 1, *P* = 0.44). There was a subtle difference between these lecithinase and β-galactosidase assays: there was variation between pherotypes in both (Fig 2B; Likelihood ratio test = 16.4, *df* = 2, *P* = 0.0003) and Group IV had the highest activity rate in 4/5 experiments, however β-galactosidase activity in Group IV and Group III was not significantly different (Likelihood ratio test = 0.83, *df* = 1, *P* = 0.36) while both Group III and Group IV pherotypes had higher levels of activity that the Group I strain (*post hoc t* = 5.69, *P* = 0.0003).

**Fig. 2.**
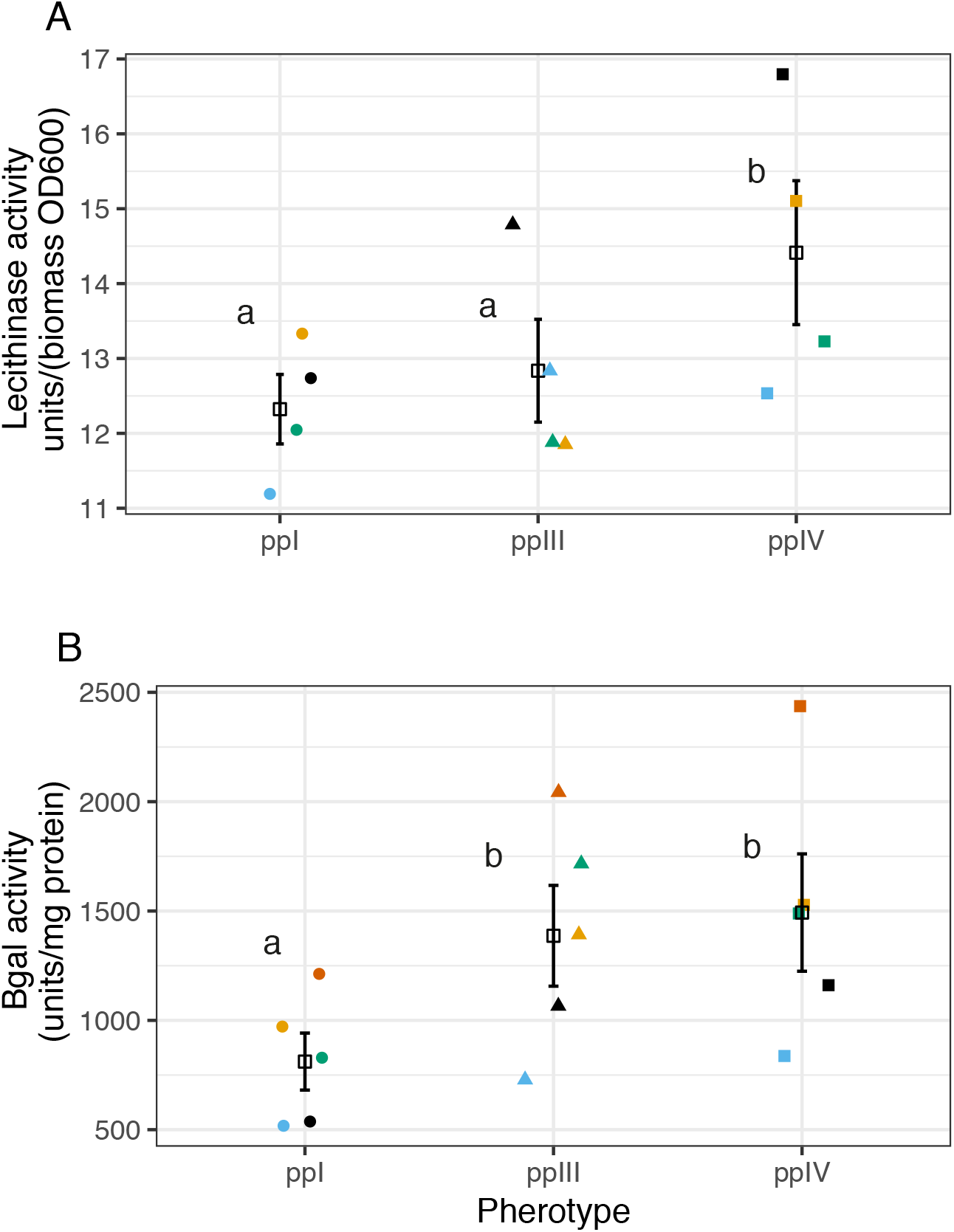
Variation in PlcR activity in near isogenic mutants with different pherotypes. This activity was determined in two different independent assays by measuring the activity of a PlcR/PapR-regulated gene product or of a PlcR/PapR-dependent promoter. (A) peak lecithinase activity-square root transformed (B) peak β-galactosidase activity. Data are independent repeats, colour coded by block, open squares and error bars are means ± SE. Different letters indicate significant differences in *post hoc* treatment contrasts. In this and all subsequent figures ppI, ppIII and ppIV indicates the Group I, III and IV pherotype mutants constructed in a common genetic background.

### Facultative cheating and productivity in homogenized host environments

Here we tested the main prediction of the facultative cheating hypothesis: that pherotypes should have a fitness advantage when rare. Based on our previous work with null mutants we further predicted that negative frequency-dependent fitness would be more likely to occur in well-mixed environments (insect homogenates) and that expression of PapR and PlcR would provide benefits in terms of increased population size (final production of spores) in insect homogenates. As predicted, there was significant variation in population size between treatment groups (Fig 3A, *F*_4,,158_ = 30.5, *P* <0.0001). The wildtype ppI strain also produced significantly more spores than the double-knockout Δ*plcR-papR* (test by model simplification *F*_1,,160_= 6.56, *P* = 0.011).

**Fig 3.**
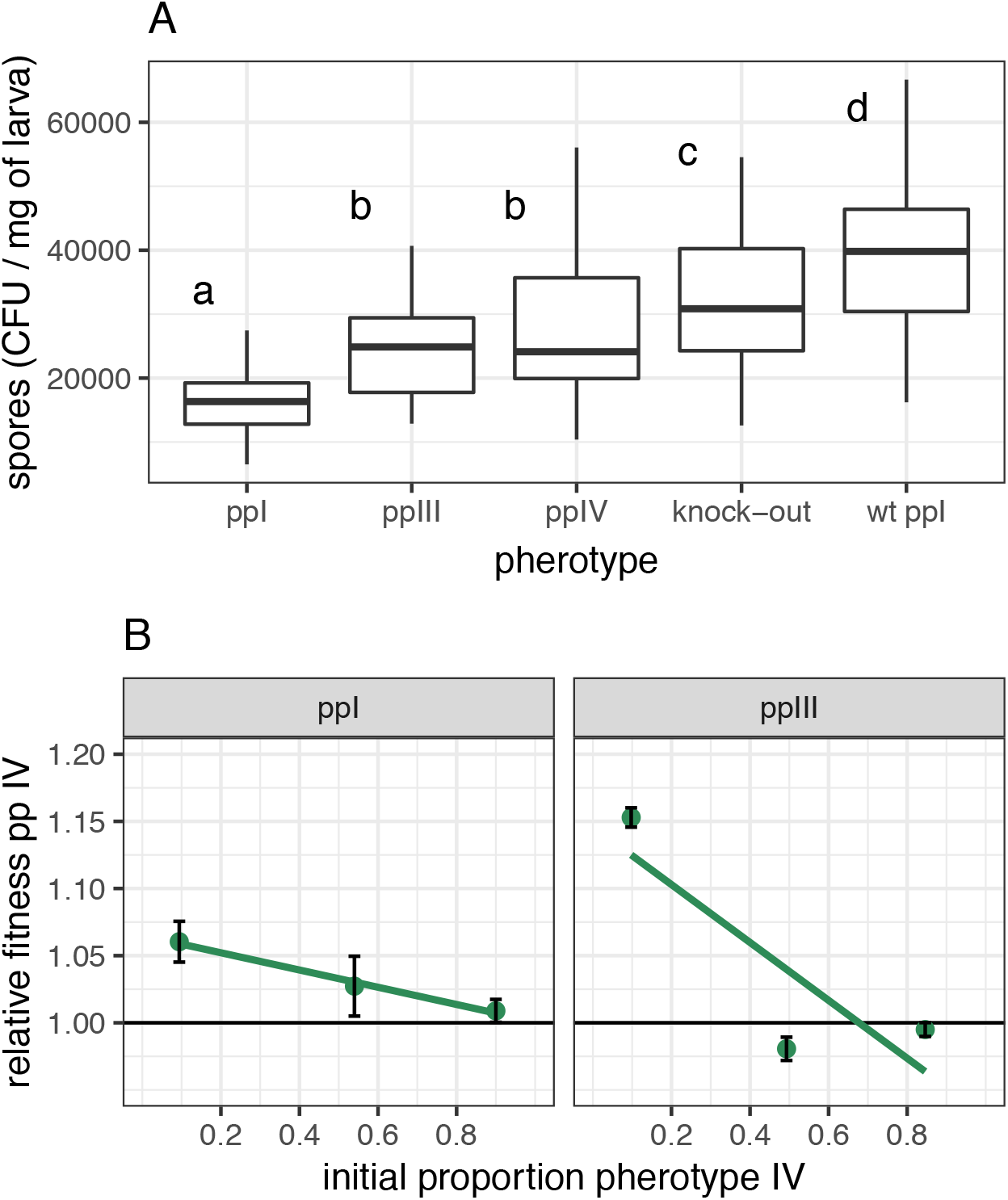
Variation in total reproduction and of relative fitness of *B. thuringiensis* pherotypes in homogenized insects. (A) Variation in final population size in numbers of spores per unit weight of insect between three pherotypes; boxplots with different letters indicate pherotypes that are significantly different in *post hoc* treatment contrasts. wt ppI indicates strain *B. thuringiensis* 407 Cry^-^ expressing chromosomally-encoded PlcR-PapR belonging to group I (B) Variation in fitness with frequency of competitors in two sets of experiments with different pherotype pairs, data are means ± SE with fitted models for each competition treatment.

We also expected that the more responsive pherotypes might have increased group level benefits and therefore produce more spores in insect homogenates. Thus while group IV and group III has similar final population size (test by model simplification *F*_1,,158_ = 0.36, *P* = 0.55); both these groups were more productive than the less responsive group I pherotype (Fig 3A, *t* = 5.48, *P* <0.0001). There was, however, a clear cost in reduced population size in the artificial plasmid construct of Group I relative to the wild type (Fig 3A, *t* = 10.4, *P* <0.0001).

As predicted by the facultative cheating hypothesis, pherotype fitness increased when rare in homogenized insect media (Fig 3B; Group IV/ III competition *F*_1,70_ = 104, *P* <<< 0.0001; Group IV / I competition *F*_1,69_ = 5.04, *P* = 0.028). Fitness patterns were not symmetrical, however. Group IV mutants, when rare, could clearly outcompete Group III (Fig 3B). However, rare Group III mutants could not outcompete Group IV (Fig 3B). Overall, the change in fitness with frequency was greatest in competitions between group IV and III (*F*_1,139_ = 17.7, *P* = 0.000045).

### Tests of facultative cheating in naturalistic infections

In naturalistic oral infections, the ecology of the experimental system is quite different and the main benefit of the PlcR system is in improved infectivity and host invasion from the midgut. In oral infections increased signal responsiveness or functioning QS does not increase population size (Fig 4A). The three pherotypes with complemented PlcR / PapR on plasmids produced fewer spores than the wild type (Fig 4A *F*_4,155_ = 11.1, *P* < 0.00001), a result consistent with a cost to using an artificial expression system, but there was no significant difference in numbers of spores produced between the Δ*plcRpapR* deletion mutant and the functional WT (Fig 4A) or between the three pherotypes (Fig 4A, *F*_2,93_ = 2.49, *P* = 0.089).

**Fig 4.**
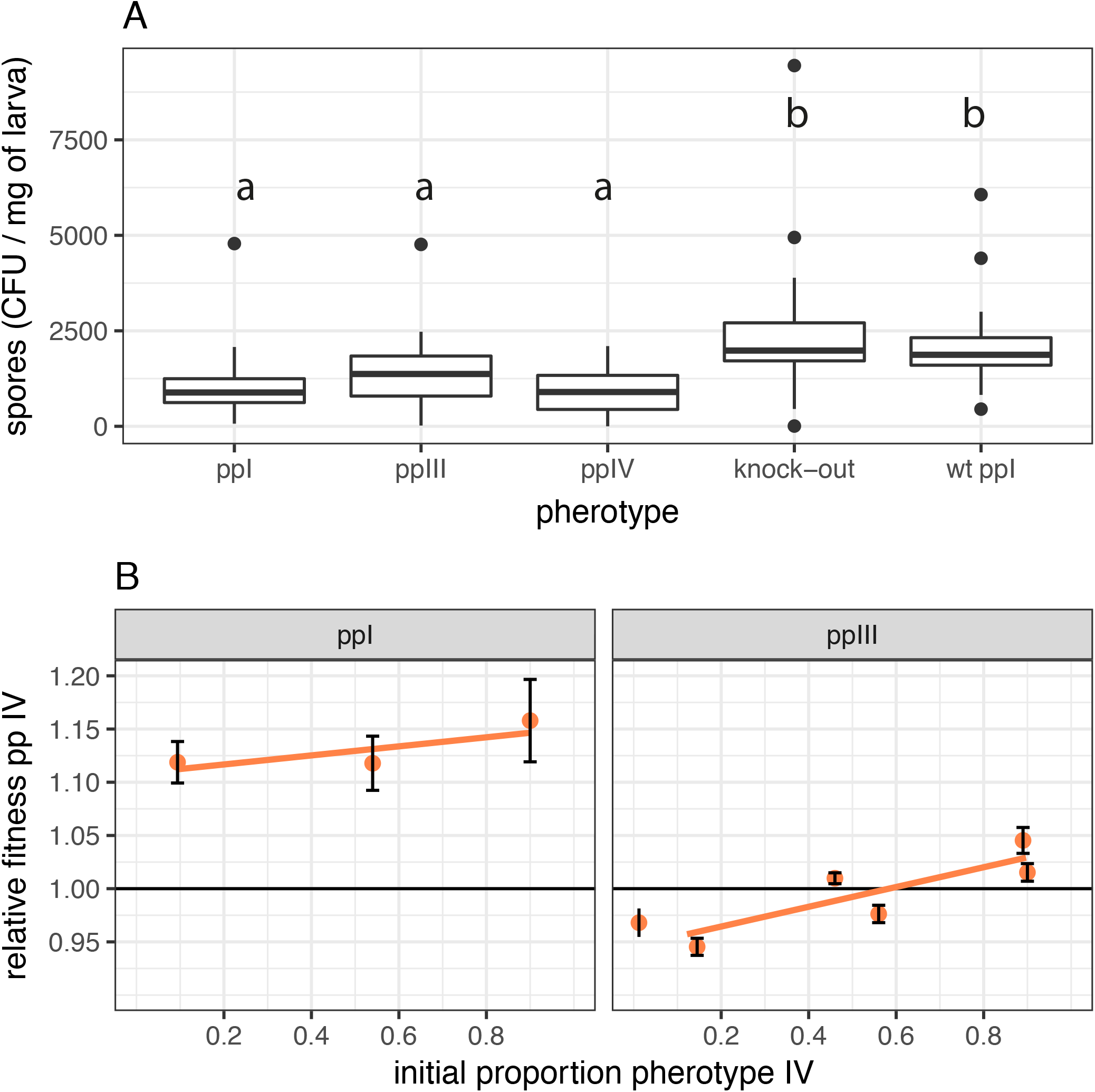
Variation in total reproduction and of relative fitness of naturalistic oral infections of insect larvae. (A) Final population size measured as spore productivity per unit weight of insect for different pherotypes, the QS knock-out null mutant Δ*papRplcR* and the QS wildtype; boxplots with different letters indicate pherotypes that are significantly different in *post hoc* treatment contrasts. (B) Variation in fitness with frequency of competitors in two sets of competition experiments with different pherotype pairs, data are means ± SE with fitted models for each competition treatment.

Contrary to the facultative cheating hypothesis, natural infections produce *positively* frequency dependent fitness in competition experiments (Fig 4B; *F*_1,587_ = 23.6, *P* < 0.0001). Note that there were clear differences in mean fitness for Group I and Group III in competition with Group IV (Fig 4B; *F*_1,587_ = 144.3, *P* < 0.0001): Group I strains had much lower fitness in this host, supporting the prediction that important phenotypic differences arise from carriage of different QS alleles. The magnitude of difference in fitness also corresponds to patterns predicted by responsiveness: the largest difference in fitness occurs between Group I and IV, where we see the greatest difference in signal responsiveness between pherotypes.

In order to validate our use of a plasmid expression system we also measured the fitness of the Group III and Group IV constructs in competition with the WT (Group I) bacterial background in naturalistic infections. Competition between Group III and Group I WT pherotypes showed positive frequency dependent fitness, (Supplementary Fig S1, *F*_1,70_ = 16.2, *P* = 0.00014), while frequency had no effect on the fitness of Group IV in competition with Group I WT (Fig S1, *F*_1,90_ = 0.07, *P* = 0.79). The relative fitness of the WT Group I strain was increased by ≈ 10% relative to that of the plasmid complemented Group I construct (Fig 3B), an effect consistent with a modest cost of plasmid carriage.

### Group size and pherotype fitness

Since QS is a density dependent process, pherotypes with quantitative differences in responsiveness to signals might have a relative fitness that varies with group size. Group size in infection is not a constant but should vary with either initial dose or the final population size of sporulated bacteria in cadavers. In the experiments with group III and IV, initial dose had a weaker effect on fitness: statistical models with final population size (AIC = −929.8) had more explanatory power than models using initial dose (AIC = −926.9). Initial doses also had no significant effect on relative fitness in addition to that of final population size in cadavers (log dose-*F*_1,425_ = 0.946, *P* = 0.33; dose * frequency interaction *F*_2,423_ = 2.55, *P* = 0.079). Dose and final population size covaried, so dose could be used as alternative explanatory variable (dose * frequency interaction *F*_2,426_ = 3.03, *P* = 0.049), in these models pherotype IV had the greatest competitive fitness at low dose and high frequencies (Supplementary Fig S2).

Since final population size was more important in determining relative fitness, we explored this variable in more detail. Pherotype IV had higher fitness in infections with lower final densities, after taking account of frequency dependence (Fig 5A, square root density * frequency interaction *F*_1,428_ = 3.61, *P* = 0.028; main effect of density *F*_1,430_ = 8.86, *P* = 0.0031). Independent experimental repeats accounted for quite substantial differences in final pathogen population size Fig 5A (*F*_1,430_ = 177, *P* < 0.0001). There are a range of explanations for the correlation between population size and relative fitness. For instance, if Group IV cells invest more in virulence and less in growth, infections dominated by Group IV might produce fewer spores.

**Fig 5.**
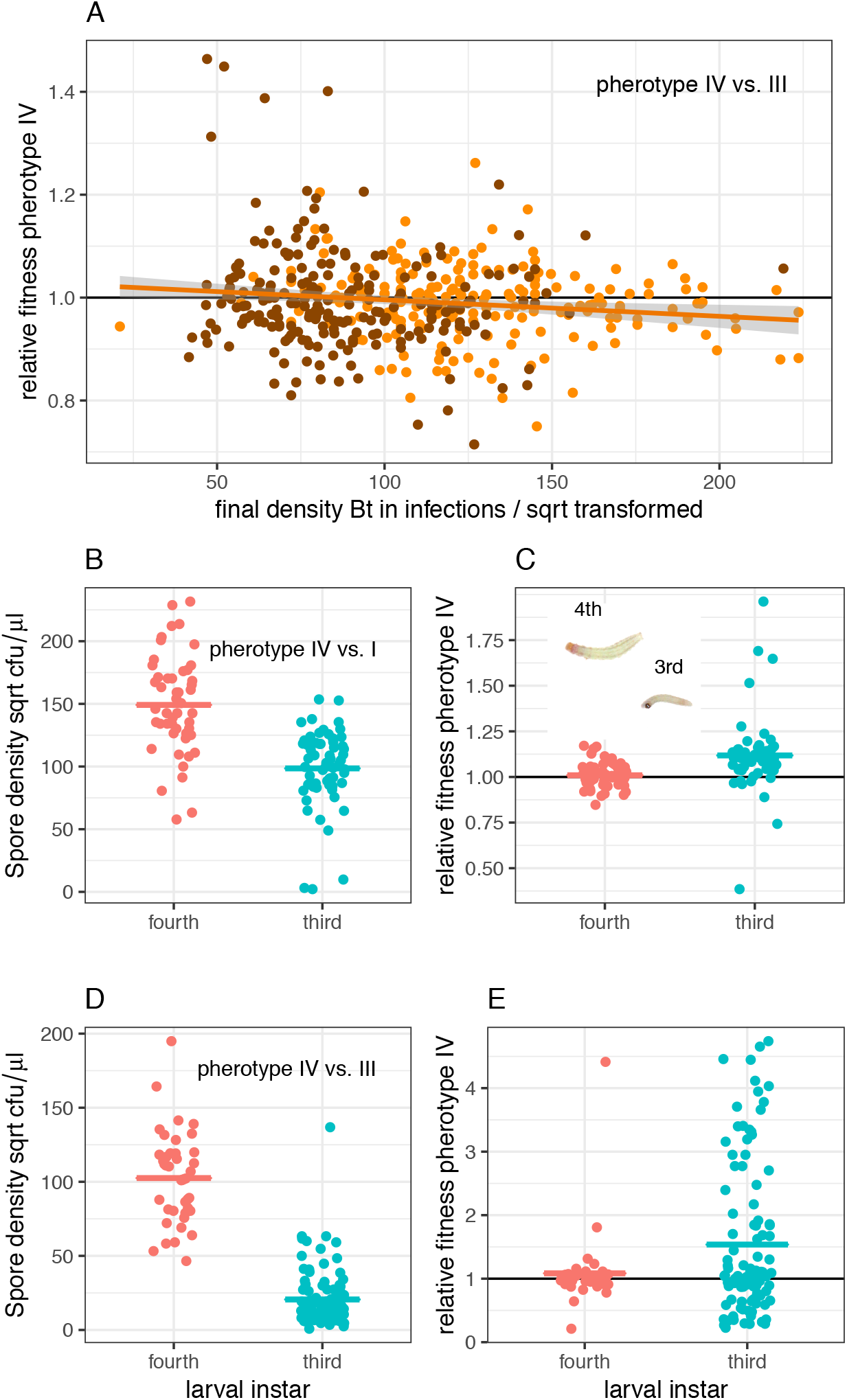
Group size and the relative fitness of competing pherotypes. (A) In competition experiments between group IV and group III, relative fitness co-varied with the final population size of *B. thuringiensis*, data are from competition experiments with 10%, 50% and 90% Group IV in inocula, colours indicate the two experimental repeats. (B) In competition experiments between group IV and group I pherotypes, different sized insect hosts produced strong differences in bacterial group size; (C) The relationship between differences in host size and relative fitness of group IV and group I pherotypes; these experiments used 50% group IV pherotype in inocula; the photograph inset shows relative larval size of fourth and third instar larvae, the fourth instar larvae is approximately 5mm in length. Competition experiments in different sized insects between group IV and group III insects also produced different final bacterial population sizes (D) and relative fitness (E). Note that, in order to display biologically informative zero counts in pairwise competitions, fitness data in E have been transformed by adding minimal countable densities to both genotypes. Where fitness of group IV > 2, no group III bacteria were detected in cadavers. Data points represent individual hosts, in A the line represent the fitted linear statistical model while horizontal bars in B, C, D and E are means.

However, the simplest explanation for the variation in pathogen population size was that insect host size varied slightly between experimental repeats and this determined the resources available for bacterial growth and thus population size dependent fitness. We therefore set up additional experiments to explicitly test the relationship between insect host size and pherotype fitness without the potentially confounding influence of variation in dose or frequency. These experiments used 50:50 mixtures of spores and competed Group IV against Group I and III pherotypes. In the experiment competing Group IV and I pherotypes, larger hosts produced much larger group sizes in terms of final populations of spores (*F_1,110_* = 59.9, *P* < 0.00001; Fig 5B). This independent experiment also confirmed that smaller insects with lower population sizes increased the relative fitness of the more responsive Group IV pherotype (*F_1,110_* = 13.3, *P* =0.0004; Fig 5C). Experiments competing Group IV and Group III pherotypes produced qualitatively similar results: the final population of spores was larger in the larger insects (*F_1,142_* = 385, *P* < 0.0001; Fig 5D). The more responsive Group IV pherotype had higher mean fitness in the smaller hosts (*F_1,142_* = 5.35, *P* =0.022; Fig 5E). However, statistical models were not well behaved for this last experiment and data were not normally distributed. Nevertheless, the difference in mean fitness in the smaller instars arose because the Group III pherotype was less efficient at invading third instar insects: in smaller hosts the Group III competitor produced significantly more zero counts than group IV (23 out of 102 cadavers versus 9 out of 102 cadavers, Fisher’s exact test *p* = 0.0114)(Fig 5E).

## Discussion

We found good support for the hypothesis that different QS pherotypes are maintained by environment dependent fitness [16]. Exchange and replacement of signalling pherotypes had detectable consequences for a range of life history traits including competitive fitness. Importantly, different QS alleles could increase or decrease the production of virulence factors or the transcription of QS regulated genes. This phenotypic variability is similar to the variation in timing and expression of QS dependent virulence factors in *Staphycoccus aureus agr* pherotypes [30]. In contrast to work on *S. aureus*, here we were able to link phenotypic variation to differences in pathogen fitness and life history *in vivo*. The most ecologically significant result was that more responsive pherotypes were fitter in smaller hosts in naturalistic infections. Importantly, in the *B. cereus* group, putative hosts encompass vertebrates, insects and nematodes with body sizes ranging over several orders of magnitude [35]. One biological interpretation here is that different pherotypes will be favoured in differently sized hosts or in ecological contexts with varying signal abundance or persistence.

Very different patterns were seen in different experimental systems – between using homogenized insects as growth media and between the oral infection that the natural mode of infection for *B. thuringiensis* [33]. First we consider how the data from naturalistic infections match up to theory. The fundamental mechanism of QS systems mean that expression of quorum regulated traits should increase with local population size or local signal abundance [2, 14]. The most successful explanation of this density dependence is based on social evolution theory [9, 10]. In these models the costs of traits are borne by individual pathogens, while the benefits are shared within groups, and as group size increases so should the total benefits to infecting pathogens [9, 10]. In this framework optimal expression of non-obligate virulence factors (formally the evolutionary stable strategy) increases with group size. This is because the benefits associated with being in larger groups eventually offsets the individual cost of investment [9, 10].

This density dependence only emerges for non-obligate virulence factors [9, 10] and so it is important to consider whether PlcR dependent virulence factors are obligate or non-obligate in the *B. cereus* group. For the ‘opportunistic pathogen’ *B. cereus sensu stricto*, virulence is almost entirely dependent on PlcR [36]. For the related insect pathogen *B. thuringiensis* the characteristic Cry toxins that define this species play a more dominant role in infection [37]. Nevertheless, in some hosts, such as *Galleria melonella*, loss of PlcR or PapR has a dramatic impact on mortality and infection [25, 36]. The host used in this study, *P. xylostella*, is highly sensitive to Cry toxins and PlcR null mutants can invade this insect successfully, indicating this QS system is not obligate for every host species. Nevertheless, QS null mutants suffer a substantial loss in their ability to invade the haemocoel of *P. xylostella* from the gut [12]. Thus, PlcR is not always strictly obligate, but when not obligate it still has a very large effect on infection success.

In addition do the products regulated by PlcR provide group level benefits than can be exploited by non-producers? Non-producers cannot outcompete functional PlcR strains in oral infections [12]. This study also shows that lower expressing variants do not consistently outcompete higher expressing variants, another key prediction of social evolution. For spatially structured infections, we might also expect negative frequency dependence, since rare non-producers should be better able to access benefits when producers are common [38]. While we have seen this pattern in competition experiments with non-producers [12], this was not repeated for the rare pherotypes in this study. Additionally, in Gram-positive species QS null mutants are rare in natural populations [12, 39, 40] supporting experimental data showing that QS non-producers have low fitness. In summary, there may be some shared benefits to PlcR regulated virulence factors, but these benefits are weak.

In addition, the positive frequency dependent fitness of competitors in this study implies that fitness increases with signal abundance but that the benefits of QS are not shared with other pherotypes in a coinfection. In this experimental system, PlcR / PapR facilitates host invasion and may occur at the level of the microcolony prior to host invasion [12]. In particular, this study also showed that less responsive pherotypes (Group III) can have reduced invasion relative to more responsive variants at low group size. Notably, priority effects (which clone enters the haemocoel first) strongly determine competitive interactions in *B. thuringiensis* [41]. Competitive differences between mutants, therefore, may occur via differences in the rate at which bacteria enter hosts. If increased expression of the lytic virulence factors regulated by PlcR / PapR favours more rapid host invasion, more responsive microcolonies may be much more competitive in smaller hosts, when group size is low and signal abundance limiting.

Predictions in relation to group size during infection are harder to evaluate, since it is hard to manipulate the number of bacteria within an ongoing infection and even harder to be confident that this manipulation occurs when quorum-sensing is happening. Group size in early infection might be affected by ingested dose, but here ingested dose had very weak effects on fitness, potentially because dose has only a transient impact on group size before spores have germinated [12] and therefore before PlcR/PapR dependent QS can begin. In contrast, host size and final pathogen density had robust impacts on the fitness of QS variants. Simply based on the size of the gut, smaller hosts should contain a reduced number of newly germinated spores in addition to placing a limit on final population size. With the above caveats in mind the observation that more productive genotypes are fitter in smaller groups contrasts with QS theory [9, 10].

In fact, the observation that optimal expression declines with group size corresponds to the predictions for obligate virulence factors and the “many hands make light work” analogy used in key theoretical analyses of QS [9]. In other words, optimal investment in virulence should decline with group size if a fixed amount of virulence factors are required for infection [9]. Other explanations for low group sizes favouring high expression are possible. For instance, the transit times of ingested material are shorter in the guts of smaller hosts. Earlier insect instars also moult and shed their gut lining more frequently so more responsive pherotypes may be fitter in smaller hosts because they can increase production of virulence factors more rapidly, and so invade hosts before they are expelled.

If PlcR does not conform to the public goods model of QS how do we explain the regulatory structure shaping density dependent expression of virulence? We hypothesize that PlcR / PapR primarily acts by sensing micro-colony formation and the spatial distribution of bacteria after attachment to the midgut [42], rather than mean population size in the host as a whole. It may be critical that PlcR is an autoregulator-activation of *plcR* increases production of the PlcR receptor protein [43] – so this QS system is driven by positive feedbacks which produce very rapid activation [44]. Since the PlcR regulon is primarily cellulytic, increased local access to host resources may provide another important positive feedback.

This study found some support for the hypothesis that facultative cheating can maintain diverse signalling systems in Gram-positive bacteria, at least in well-mixed environments with much reduced spatial structure. Facultative cheating requires a number of particular ecological and biological contexts, such as the close proximity of colonies that exchange signals by diffusion, and intermediate levels of bacterial population structure. Under a clonal population structure genotypes only interact with themselves, so there is no opportunity for cheating. The high levels of clonality in *B. thuringiensis* and *S. aureus* infections make facultative cheating more unlikely [13, 41, 45]. Conversely, if there is too much mixing of genotypes, then theory predicts that null QS cheaters will predominate and QS systems will collapse [28, 32]. Another important assumption is that there must be low level of cross-talk between different pherotypes, i.e. weak activation of receptors by signal from different groups. The limited level of cross-talk between the *B. cereus* group pherotypes did not preclude facultative cheating, provided there was reduced spatial structure. There were also asymmetries in fitness for rare variants that could be nicely explained by asymmetries in cross talk. For example the group IV pherotype, when rare, could successfully exploit group III when common, but the reverse was not true. This was potentially because group IV responds very weakly to the signals of group III and therefore should be a better facultative cheat.

In previous evolutionary studies of QS, signalling and QS-regulated traits are public goods, with strong indirect fitness benefits and high fitness at low frequencies [4, 7, 9, 10, 46], patterns that only hold in insect homogenates in this study. However, PlcR does not conform well to a public goods model in insect infections: group level benefits are very weak and the variation in fitness with group size corresponds to predictions for obligate virulence factors, rather than for typical QS traits. We therefore need an alternative explanation for the density dependent regulation of virulence for PlcR. Social evolution models cannot also easily explain the observed polymorphism in peptide signals. A model of environment dependent fitness for QS variants seems more relevant for some of the best studied Gram positive pathogens: *S. aureus* and the *B. cereus* group, since different allelic variants can show differences in gene expression, fitness, life history and potentially distinct host niches [47, 48].

## Materials & Methods

### Bacterial Strains

Distinct *plcR-papR* genes were sourced from three of the four pherotype groups found in *B. cereus sensu lato*. Group IV, Group III and Group I genes were derived from *B. thuringiensis* serovar *roskildiensis, B. thuringiensis* subsp. *kurstaki* and *B. thuringiensis subsp berliner* respectively and introduced on the stable pHT304 plasmid [49] into the acrystalliferous strain *B. thuringiensis* 407 Cry^-^ A’Z Δ*plcR-papR* which contains a *lacZ* fusion for reporting *plcR* transcriptional activity [23]. Experiments also used the 407 Cry^-^ A’Z Δ*plcRpapR* mutant lacking the complemented plasmid as well as the wild-type *B. thuringiensis* 407 Cry^-^ isolate with the intact chromosomal *plcR-papR* genes, which was the source of the pherotype Group I gene, produced as described previously [25]. In order to distinguish between pherotypes, the Group IV plasmid strain was constructed to be tetracycline-resistant while the Group I and Group III plasmids retained the erythromycin-resistance gene as a marker. Details of cloning procedures are described fully in the Supplementary Information (SI).

### PlcR activity assays

Two assays were used to compare the expression of genes under control of the PlcR QS system in standard laboratory conditions. The first assay assessed the hydrolysis of egg yolk phosphatidylcholine by the lecithinase PlcB [50], whose encoding gene is a member of the PlcR regulon in *B. cereus* [51]. In the second set of assays, we measured the transcription of the *plcA* gene, also under the control of PlcR [43], using a transcriptional fusion between the promoter of this gene and the reporter gene *lacZ* encoding a ß-galactosidase. Specific activity in the ß-galactosidase and lecithinase assays was measured as described previously [50] and repeated five and four times respectively. Lecithinase activity is calculated as the slope of the linear part of the curve/[OD600 of the culture at the time of sampling × volume of supernatant used in the assay (L)], while ß-galactosidase is calculated according to (OD420 x 1500000)/(reaction time in min x vol crude extract in μL x C) where C is the protein concentration.

### Infection experiments

Competition experiments using oral infection of diamondback moth *Plutella xylostella* larvae or culture in homogenized insect medium followed published protocols [12], except that insect homogenates were not pasteurized before maceration. All experiments with homogenates used shaken cultures in 24 well plates containing two sterile glass beads. All larvae were reared aseptically on sterile diet after being hatched from surface sterilized eggs [52]. Competition experiments based on oral infections of Group III and Group IV pherotypes used a full factorial design with 5 frequencies of competing strains (approximately 100%, 90%, 50%, 10% and 0% Group IV) and three doses (2×10^3^, 1×10^4^ and 5×10^4^ spores μl^−1^) with 25 insects (third instar) in each treatment. These experiments were repeated twice. Competition experiments using Group I bacteria used three frequencies (90%, 50%, 10% Group I) and a dose of 1×10^4^ spores ^−1^ and 30-35 larvae per treatment. Competition experiments in hosts of different size used assay used third and fourth instar *P. xylostella* larvae with 50:50 mixtures of Group I and IV pherotypes and a 50:50 mix of Group III and IV pherotypes.

### Data analysis

Bacterial relative fitness (W) was calculated using the ratio of Malthusian parameters or relative growth rate (*W_A,B_* = *log2* ((final^A^/initial^A^))/ *log2* ((final^B^/initial^B^)) where final^A^ and final^B^ *etc*. refer to the final and initial numbers of strains A and B [53]. In order to calculate *W* we estimated the initial total population of bacterial founders in each infection at 50 cells. This is a high estimate based on previous experiments [41], although relative fitness is not particularly sensitive to this parameter. It is not possible to calculate relative fitness when there is no detectable growth in hosts from either competitive partner. In the Group III & IV competition experiments with different sized larvae, zero cell counts were transformed by addition of the minimal detectable value, principally to allow visualization of results. Other data were log10 or square root transformed to improve homoscedasticity as required. Relative fitness and bacterial reproduction were analysed using generalized linear modelling (glms); peak enzyme activity data were analysed in mixed linear effect models with experimental replicate as a random factor. Planned *post hoc* comparison in glms were conducted using model simplification and pooling of treatment levels. Where error structures or data failed to meet tests of normality or homoscedasticity (produced by zero counts) data were analysed with classical tests (Fisher’s exact test). All statistical analyses were performed in *R v3.3.2* (R Core Team 2013).

### Data availability

Experimental data on expression, bacteria densities and relative fitness supporting this publication are openly available from the University of Exeter’s institutional repository at: https://doi.org/10.24378/exe.XXX. [This is a a placeholder doi, unique doi to be allocated following acceptance.]

## Acknowledgements

This work was supported by an NERC Research Fellowship award (to BR), and by the MICA department of INRA (National Institute of Agronomical Research). Thanks to Lucy Scott and Molly Raymond for technical support.

## Supplementary Information

A single file containing supplementary materials and methods; supplementary figure S1 and supplementary figure S2.

